# SimMS: A GPU-Accelerated Cosine Similarity implementation for Tandem Mass Spectrometry

**DOI:** 10.1101/2024.07.24.605006

**Authors:** Tornike Onoprishvili, Jui-Hung Yuan, Kamen Petrov, Vijay Ingalalli, Lila Khederlarian, Niklas Leuchtenmuller, Sona Chandra, Aurelien Duarte, Andreas Bender, Yoann Gloaguen

## Abstract

Untargeted metabolomics involves a large-scale comparison of the fragmentation pattern of a mass spectrum against a database containing known spectra. Given the number of comparisons involved, this step can be time-consuming. In this work, we present a GPU-accelerated cosine similarity implementation for Tandem Mass Spectrometry (MS) with approximately 1000-fold speedup compared to the MatchMS reference at a rate of 0.005% incorrect matches and a rate of 0.002% incorrect scores. We describe the underlying reasons for these errors and provide means to avoid them.

## Introduction

In the field of untargeted metabolomics, Tandem Mass Spectrometry (MS/MS)^1^ is a well-established technique for identifying compounds within complex biological samples. The process works by comparing an unknown compound’s mass spectrum fragmentation pattern (“query”) against a database of known spectra (“reference”) in an effort to identify the unknown compound’s chemical composition^2^.

Cosine similarity and its variants are popular methods facilitating MS/MS spectra comparison^3,4^. Cosine similarity works by calculating a cosine of the angle between vectors of fragmentation intensities (“peaks”). Since an exact match between two fragmentation spectra isn’t practically possible due to measurement errors, finding the cosine score involves solving an *assignment problem* between the two sets of spectral peaks, with the goal of finding a valid matching of peaks (within *m/z* tolerance) that maximize the value of the cosine. Calculating the exact cosine similarity (Hungarian cosine^5^) is usually impractical for even moderate numbers of spectra. “Greedy” cosine similarity is an efficient approximation of the cosine similarity, which solves the assignment problem using a greedy heuristic^6^. “Modified” cosine similarity is an extension of greedy cosine that uses the precursor mass as an additional input and has been shown to outperform greedy cosine in some cases^3^.

Huber *et al*. also introduced MatchMS^6^, an open-source MS/MS processing library in python. This library allows implementing easy-to-reproduce workflows to process raw mass spectral data into more useful forms (i.e. molecular networks). MatchMS conveniently implements all three types of cosine similarities. Unfortunately, while the MatchMS implementation is convenient, it is too slow for processing spectra on the scale of 10^10^ pairwise comparisons or more. At such scales, which are routine in our untargeted metabolomics workflows, MatchMS requires tens of CPU-days, necessitating a search for more efficient and robust approaches^7-9^.

To address this issue, Harwood *et al*. introduced BLINK^10^, which approximates the greedy cosine by first blurring the query spectral peaks, transforming it into a sparse matrix, and then performing sparse matrix multiplication between the query matrix and the reference matrix to directly compute the score matrix. This effectively side steps solving the peak assignment problem, allowing BLINK significant speed improvements over MatchMS. Unfortunately, BLINK rapidly loses accuracy when tolerance is larger than 10^−2^.

In the current report, we speed up Cosine calculation from an engineering point of view. We leverage the fact that cosine calculation is readily parallelizable and rewrite both greedy and modified cosine algorithms using CUDA^11^ into a single, highly optimized GPU kernel that, depending on underlying GPU hardware, can process spectra up to 1,000 times faster than MatchMS. We show that the approach exactly replicates MatchMS results and supports a much wider tolerance range without compromising accuracy. Further, we distribute the kernel into a user-friendly Python package that can act as a drop-in replacement for respective MatchMS cosine similarity classes. All code, results, and notebooks supporting this report are freely available under the MIT license at https://github.com/pangeAI/simms/.

## Results

We found that using the Cosine kernel implementation presented in this work with a single NVIDIA A100 GPU for calculations^12^ is approximately 1,000x faster than using MatchMS for both greedy and modified cosine similarities, as shown in Figure 1. Calculating greedy cosine similarity at the scale of 100,000 queries paired with 1.5 million reference spectra takes an estimated 13 weeks with MatchMS, in contrast to only 3 hours using the kernel.

**Figure 1:**
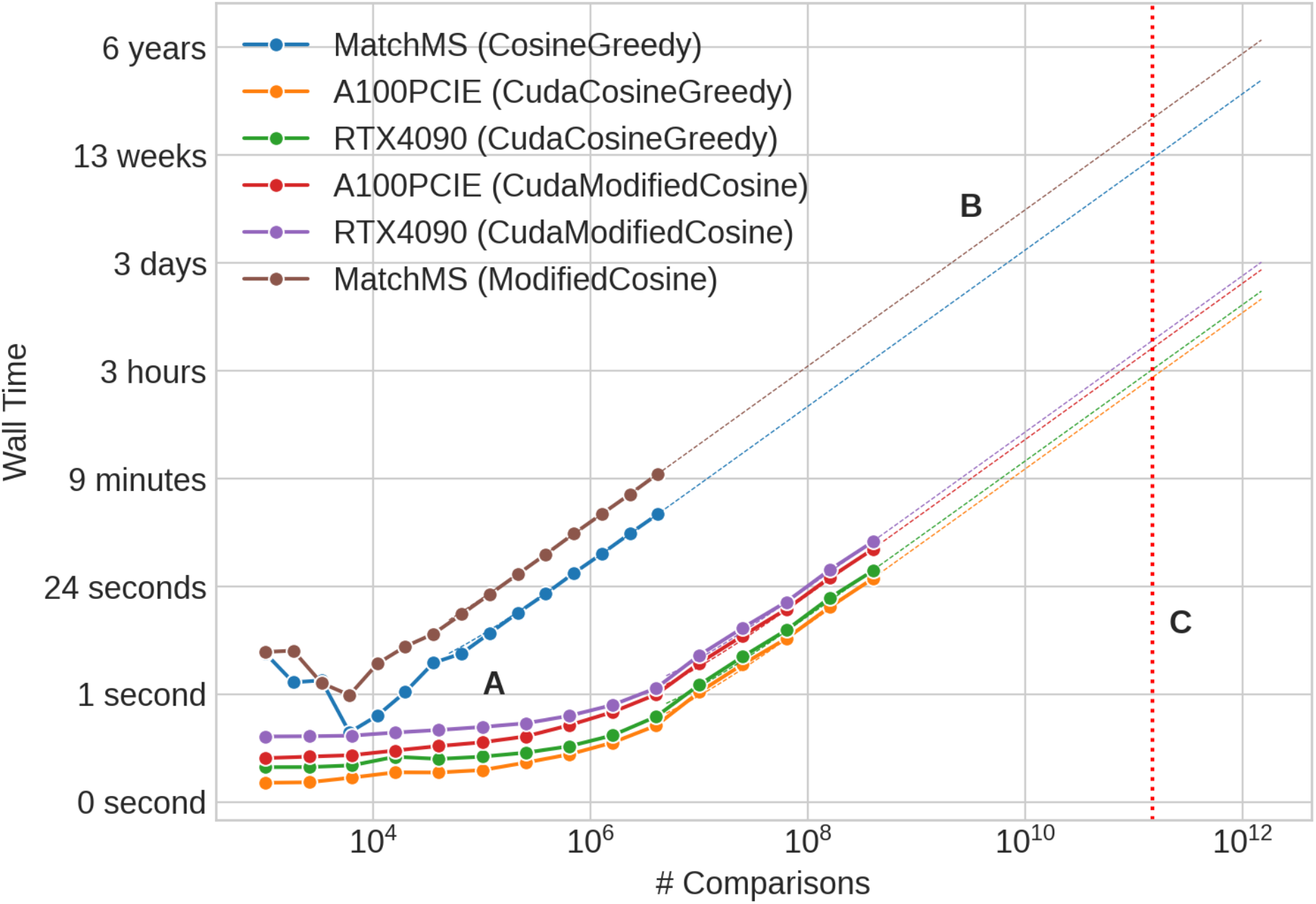
Cosine scoring runtimes for GNPS subsets with default parameters, with different GPUs and methods. For example, the red line shows the SimMS performance on A100 for modified cosine scoring, and the brown line is the same calculation with MatchMS. Marker dots (A) represent real measurements, and dashed lines (B) are their linear interpolations. (C) is a marker of 1.5·10^11^ comparisons, and represents our target processing goals. Notably, all approaches seem to have a similar asymptotic computational complexity.

Modified cosine similarity is slower than greedy cosine (also shown in Figure 1), independent of hardware used, since the steps required to calculate the modified cosine score includes all steps required for calculating the greedy cosine score^3^.

We performed a direct comparison of results for predicted scores and the number of matches, as shown in Figure 2. We found that for default parameters, the kernel and MatchMS results are within ±0.001 of each other 99.99% of the time. Since the kernel is algorithmically equivalent to its MatchMS counterpart, the few errors stem from peaks that are almost exactly tolerance distance apart. In other words, when paired with MatchMS, these peaks appear to be inside tolerance distance, but processing them with a GPU changes their binary floating point representation, which is enough to make them appear outside of the tolerance distance. When a spectrum has a single very intense peak, such binary representation changes can result in large score errors. The score comparison plots in panels Figure 2A and Figure 2C show these kinds of errors in the bottom right.

**Figure 2:**
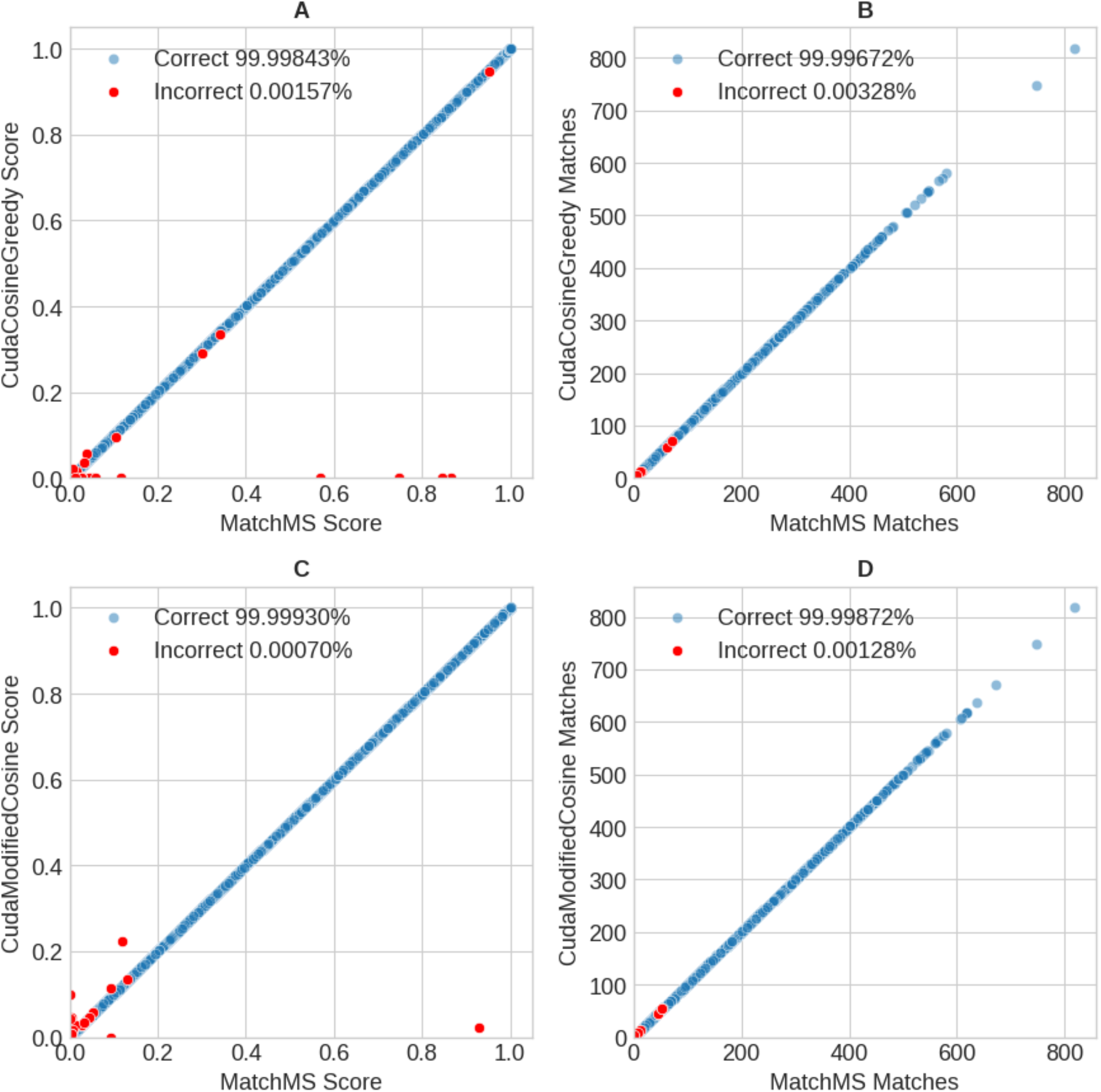
Direct comparison of scores of 4.1M random GNPS spectra pairs, illustrating SimMS error patterns. Red dots denote incorrect (absolute error > 0.001) SimMS outputs. (A) and (B) show greedy cosine results between SimMS and MatchMS, performed at float32 precision. In (C) and (D) we compare modified cosine results at float64 precision. Occasional large errors in (A) and (C) are caused by spectra with very few (one or two) large peaks.

The rate of incorrect scores is usually lower than the rate of incorrect number of matches. This asymmetry originates from the filtering step. Filtering usually removes most of the low-intensity matches from the score calculation, thus making the missing or extra match unlikely to affect the score value.

In Figure 3 we can see how the changing tolerance and match limit influence SimMS performance. In Figure 3A and 3D we see that lowering these values can significantly speed up spectra processing time, but this comes at the price of increasing the overflow rate, as seen in Figure 3B and 3E. The accuracy rate in Figure 3C and 3F is approximately equal to the 1 - overflow rate.

**Figure 3:**
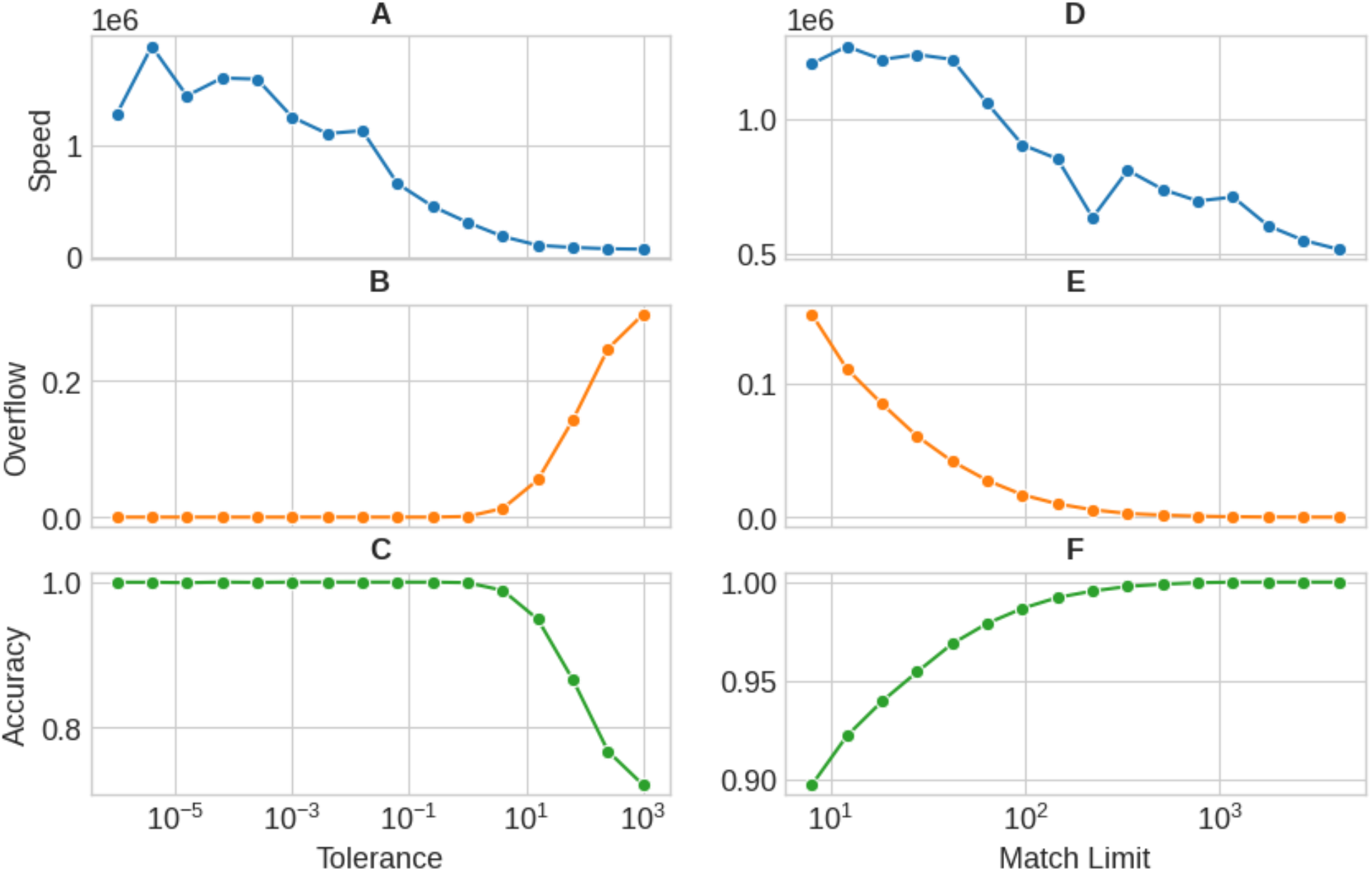
Key metrics as a function of changing tolerance (A) - (C) and match limit (D) - (F). In a and d speed means the number of comparisons per second. “Overflow” is the proportion of scores that are returned with the overflow flag. Accuracy is the proportion of scores that are within ±0.001 of MatchMS.

## Methods

### Implementation

We used as input a list of references and queries, denoting their respective lengths as **R** and **Q**. For both lists, consecutive spectra are grouped into batches of size **B**, the last batch contains leftover spectra and padding, as needed.

Inside each batch, all spectra are concatenated into a single ℝ^2, *B, M*^ tensor, where **M** is the number of peaks in the longest spectrum in that batch, **B** is the batch size (Figure 4a). A batch contains stacked peak m/z and intensity values in the first dimension. Spectra that are shorter than **M** are padded with zeros. Spectra that are larger than **M** are truncated with an argument *N*_*max peaks*_.

**Figure 4:**
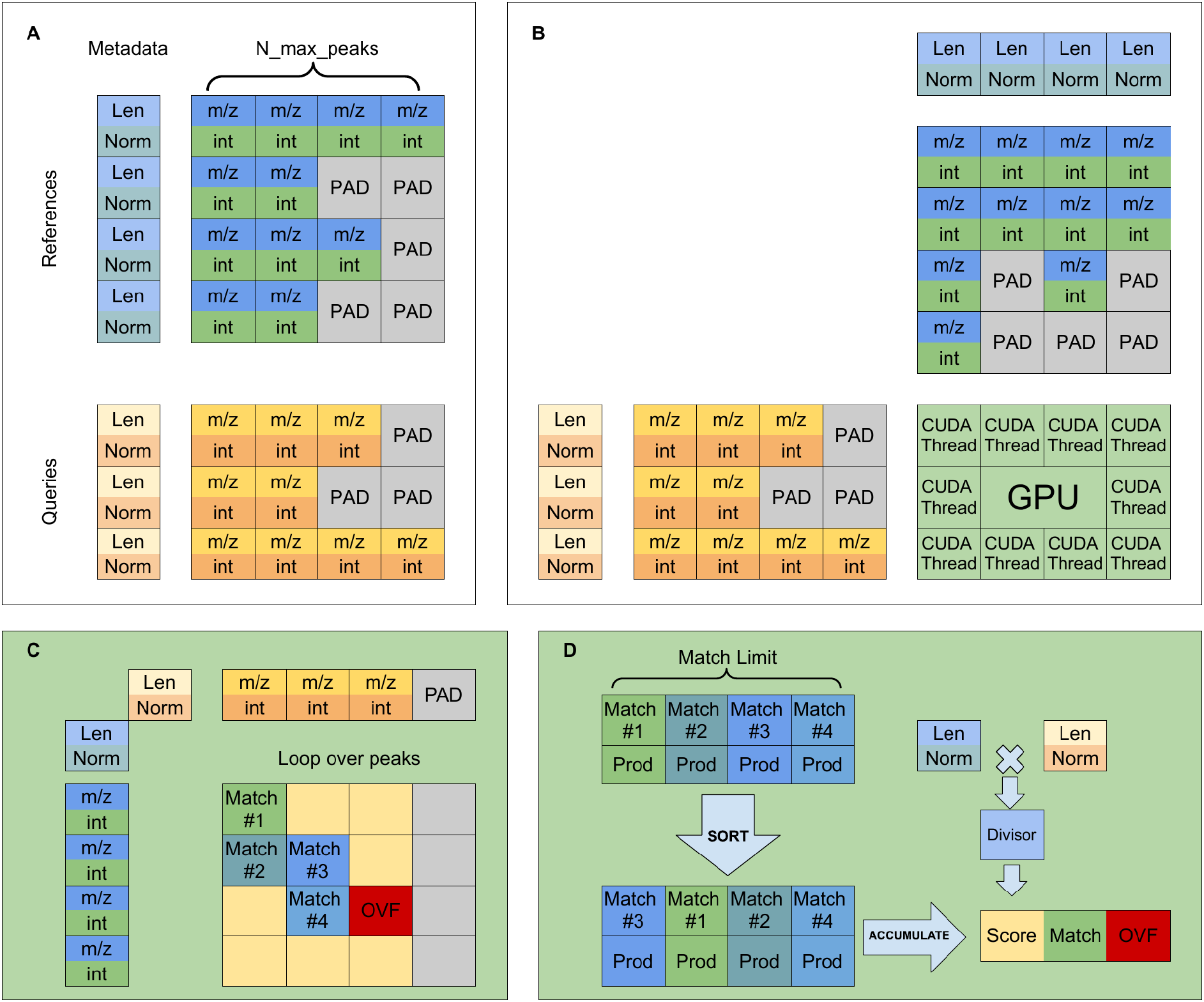
Overview of processing on GPU. In a first step (A), we pack all spectra and metadata in 3D and 2D tensors, respectively. In a second step (B), we align the spectra with the GPU grid. Steps (C) and (D) happen within a single GPU thread. In (C), we accumulate potential matching peaks up to the given match limit. In the last step (D), we sort, deduplicate and reduce matched peaks, returning three values.

The “metadata” tensor is created alongside the spectra batch. The metadata tensor contains the length, cosine norm, and, in case of modified cosine, the precursor m/z values. Metadata tensor is a ℝ^*K,B*^ tensor, where K is either 4 or 6, and **B** is the batch size. For greedy cosine, dimension **K** is 4 and consists of lengths of reference and query spectra and norms of reference and query spectra. In case of modified cosine metadata additionally contains precursor m/z values for reference and query spectra. For batches corresponding to leftover spectra, we pad the resulting empty space with zeros.

The full similarity matrix of size *R* × *Q* is infeasible to store in GPU memory. Therefore processing is done block-by-block. The full *R* × *Q* grid is split into *B* × *B* sized, non-overlapping blocks. In total this results in 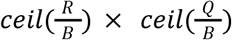 number of blocks. For each block an output ℝ^3,*B,B*^ tensor is allocated on the GPU. A pair of batches of spectra, and their respective metadata tensors are transferred to the GPU. The tolerance, m/z power, intensity power, *N*_*match limit*_ and *N*_*max peaks*_ are supplied to the kernel as compilation-time constants. At this point the kernel is launched.

The kernel is written using Numba^13^. Inside the kernel, a single CUDA thread is assigned to calculate the cosine score between one reference spectrum and one query spectrum from the supplied batches. The computing power of SimMS stems from the fact that modern GPUs can process tens of thousands of threads simultaneously, allowing us to process each block in a fraction of a second.

The kernel itself consists of three main stages (Figure 4C and 4D). First, pairs of peaks within tolerance are collected. A maximum of *N*_*match limit*_ pairs are collected. If the number of pairs exceeds this limit, an overflow flag is raised and the collection stops early. Next, the pairs are sorted by the value of the product of the intensities of paired peaks. Finally, the ordered pairs are deduplicated and accumulated into an unnormalized cosine score. An auxiliary boolean array is used to mark each accumulated peak, to avoid duplicate contributions to the score. As a final step, the two norms from metadata are multiplied to obtain the normalizing constant, which divides the unnormalized score to produce the final cosine score (Figure 4D).

After the execution, the block output tensor contains 3 results for every reference/query pair in the batch. These are: score, number of matches, and overflow (binary flag). Each block is then concatenated together in order to form the full *R* × *Q* similarity matrix. In case of processing a very large set of references and queries (larger than 100,000), the required memory to store the full similarity matrix as a dense array is impractically large. Additionally, we find that most of the scores are lower than 0.5. For such cases we use “sparse” implementation, where the similarity matrix is filtered to discard all results with score below a user-defined “sparse threshold” and then store the remaining entries as a sparse matrix in DOK format.

Finally, we concatenate all the results and return them, either as a dense array or as a sparse DOK matrix.

### Hardware

All of our kernel and MatchMS experiments were performed on rented Vastai (Vast.ai) instances. Preferably, we rented instances with at least 16GB RAM, 8 or more CPUs, and at least a single RTX4090 GPU. Performance of the original MatchMS algorithm is independent from GPU, while our kernel performance heavily depends on it - A100 GPU usually outperforms RTX4090, which in turn outperforms older GPUs.

### Interoperability

We have taken care to make the SimMS package fully compatible with MatchMS. During the implementation of the kernel, we also discovered that Cosine Greedy scores from MatchMS were semi-randomly fluctuating when run on different CPU hardware, causing the exact MatchMS results to be unrepeatable. We later patched this bug, and from version 0.24.0 onwards, the patch has been merged into the core MatchMS package. Correction details are available at https://github.com/matchms/matchms/pull/596.

The full code is available on GitHub at https://github.com/PangeAI/simms under an MIT license.

