## Supplementary information for "SimMS: A GPU-Accelerated Cosine Similarity implementation for Tandem Mass Spectrometry"

### The kernel algorithm

Python

```
def kernel(reference, query, info) -> float, int, bool:
        matches = array(max_size=match_limit)
        score_norm = get_rnorm(info) * get_qnorm(info)

        # 1. Collect peaks
        for r, q in cartesian(reference, query):
                if matches.has_space():
                        if abs(r - q) < tol:
                                matches.append((r, q))
                else:
                        overflow = 1; break
        # 2. Sort peaks
        matches = sorted(matches)

        # 3. Filter
        visited = array(max_size=n_max_peaks, fill=False)
        score, num_matches = 0, 0
        for r, q in matches:
                if not (visited(r) or visited(q)):
                        score += get_peak_product(r, q)
                        num_matches += 1
                        visited[r], visited[q] = True, True

        score = score/score_norm
        return score, num_matches, overflow
```

**Algorithm S1. Kernel pseudo-code.**
